# CRISPR screens and lectin microarrays identify novel high mannose N-glycan regulators

**DOI:** 10.1101/2023.10.23.563662

**Authors:** C Kimberly Tsui, Nicholas Twells, Emma Doan, Jacqueline Woo, Noosha Khosrojerdi, Janiya Brooks, Ayodeji Kulepa, Brant Webster, Lara K Mahal, Andrew Dillin

## Abstract

Glycans play critical roles in cellular signaling and function. Unlike proteins, glycan structures are not templated from genes but the concerted activity of many genes, making them historically challenging to study. Here, we present a strategy that utilizes pooled CRISPR screens and lectin microarrays to uncover and characterize regulators of cell surface glycosylation. We applied this approach to study the regulation of high mannose glycans – the starting structure of all asparagine(N)-linked-glycans. We used CRISPR screens to uncover the expanded network of genes controlling high mannose surface levels, followed by lectin microarrays to fully measure the complex effect of select regulators on glycosylation globally. Through this, we elucidated how two novel high mannose regulators – TM9SF3 and the CCC complex – control complex N-glycosylation via regulating Golgi morphology and function. Notably, this method allowed us to interrogate Golgi function in-depth and reveal that similar disruption to Golgi morphology can lead to drastically different glycosylation outcomes. Collectively, this work demonstrates a generalizable approach for systematically dissecting the regulatory network underlying glycosylation.

## Introduction

All living cells and organisms are covered with glycans – complex carbohydrates linked to proteins, lipids, and RNA^1^. Glycans play critical roles in many biological processes – intracellularly, glycans are essential for protein folding and influence the stability, localization, and activity of many proteins within and outside the cell. Extracellularly, glycans on the cell surface mediate cell-cell recognition and interactions, including many immunological responses^2^. Under many acute and chronic disease states, glycosylation can become dysregulated and actively contribute to disease progression^34–6^. For example, the high mannose glycan epitope, typically found intracellularly within the ER and the Golgi, was recently identified as a stress signal for influenza virus infection, and their cell surface presentation is suggested to cause excessive tissue damage through binding innate immune lectins and over-activating the complement pathway^78^. However, how high mannose or other glycan motifs are regulated at the cell surface remains relatively unknown.

Understanding how cell surface glycosylation is regulated has been historically challenging due to the non-templated nature of glycans. Unlike proteins, biosynthesis of glycan structures is not directly encoded in genes. Instead, glycan synthesis is controlled by an expanded network of genes that regulate biosynthetic enzyme expression and localization, glycan trafficking, organelle function, substrate availability, and carbohydrate metabolism, producing a heterogeneous collection of glycans on the cell surface^9,10^. While the biosynthetic enzymes that directly catalyze glycosidic linkages are mostly mapped out through decades of dedicated research^10,11^, the contribution of other genes remains relatively poorly understood. In addition, changes to cell states, such as activation of proteostasis stress response pathways^12^, can also drastically alter the glycan repertoire of a cell, adding to the difficulty of understanding how glycosylation is regulated.

While understanding the biology underlying a specific glycan epitope remains challenging, recent advances in glycomic techniques have enabled a comprehensive survey of the glycan landscape of cells and tissues under healthy and disease states^13^. In particular, lectin microarrays, which utilize a variety of lectins and antibodies to detect specific glycan moieties, have proven to be a powerful and highly sensitive method for uncovering glycosylation differences between biological samples^14^. Lectin microarray analyses have identified glycan changes across many diseases and have been useful for biomarker discovery in predicting disease outcome and vaccine response^7,15–18^. However, lectin microarrays alone cannot readily reveal the underlying biology that causes the glycan change in the first place.

Recent advances in CRISPR screening have proven to be a powerful tool for understanding the genetics of glycosylation. Utilizing bacterial and plant toxins that bind known glycan moieties, novel genetic regulators have been identified that control the synthesis of glycoproteins and glycolipids ^19–21^. Expanding on these works, we utilized the accumulated knowledge of naturally isolated lectins and their binding specificities in CRISPR screens and lectin microarrays to identify and characterize novel regulators of cell surface N-glycosylation. Specifically, we applied our strategy to uncover regulators of high mannose glycans – the essential intermediate structure for all N-glycans and an important glycan epitope of the innate immune response.

We first used FACS- and magnetic-based cell sorting methods to conduct genome-wide and targeted screens to uncover the expanded network of genes that control cell surface levels of high mannose N-glycans. Next, we employed lectin microarrays to measure the glycan changes comprehensively to obtain mechanistic insights into how select regulators control cell surface glycosylation. Through this, we discovered how two novel regulators of high mannose glycosylation – a previously poorly characterized gene TM9SF3 and the endocytic recycling machinery CCC complex – control complex N-glycosylation and Golgi morphology and function. Specifically, we found that loss of TM9SF3 function reduces cis- and trans-Golgi colocalization and inhibits complex N-glycan formation, while disruption to the CCC complex leads to Golgi fragmentation yet mildly increases cis/trans-colocalization, enhancing complex N-glycan production. Notably, the unbiased interrogation of Golgi function using lectin microarray revealed that Golgi morphology changes that are similar on a surface level (i.e. fragmentation) can lead to drastically different glycosylation outcomes. Together, these findings reveal novel cell surface high mannose N-glycosylation regulators and validate the strategy to combine CRISPR screening with lectin microarray technologies for revealing novel regulators of glycosylation.

## Results

### UPR^ER^ activation upregulates high mannose glycans on and within cells

All N-glycans begin as the 14-sugar glycan structure (Glc_3_Man_9_GlcNAc_2_), which is added onto selected asparagine residues on nascent proteins as they enter the ER for folding^22^. As these glycoproteins mature through the ER and cis-Golgi, they transition through a high mannose stage (Man_5_-Man_9_) after initial processing steps that trim off glucose residues. Typically, these high mannose structures are further processed into more complex glycans in the Golgi, such as those elongated with repeating units of Gal and GlcNAc (“poly LacNAc”) or capped with sialic acid residues, resulting in an extensive array of mature, complex N-glycans at the cell surface^22^. When cells experience stresses such as influenza viral infection, high mannose glycans can become upregulated at the cell surface and function as a stress signal that binds innate immune lectins^7^. However, it is unknown how healthy cells maintain low levels of high mannose at the surface or how these glycans become upregulated under stress conditions. Thus we chose to focus on the regulation of high mannose glycans with our approach.

Unfolded protein response (UPR) activation through the XBP1 pathway was required for high mannose expression in the human lung carcinoma cell line A549 in response to influenza and, in other work, was shown to induce high mannose in other cell lines in a cell-type specific manner^7,12^. We thus focused on understanding how high mannose glycans are regulated under basal and UPR^ER^-activated conditions. To this end, we built a doxycycline(dox)-inducible system to enable the overexpression of XBP1s, the key transcription factor that mediates IRE1 branch of the UPR^ER^ response (**Fig. 1a**). This dox-inducible XBP1s system was lentivirally introduced into the cell line A549s. Consistent with previous work that used similar strategies to activate branches of the UPR^ER 12,23^, our system also enabled specific upregulation of XBP1s targets (**Fig. 1b**).

**Fig. 1:**
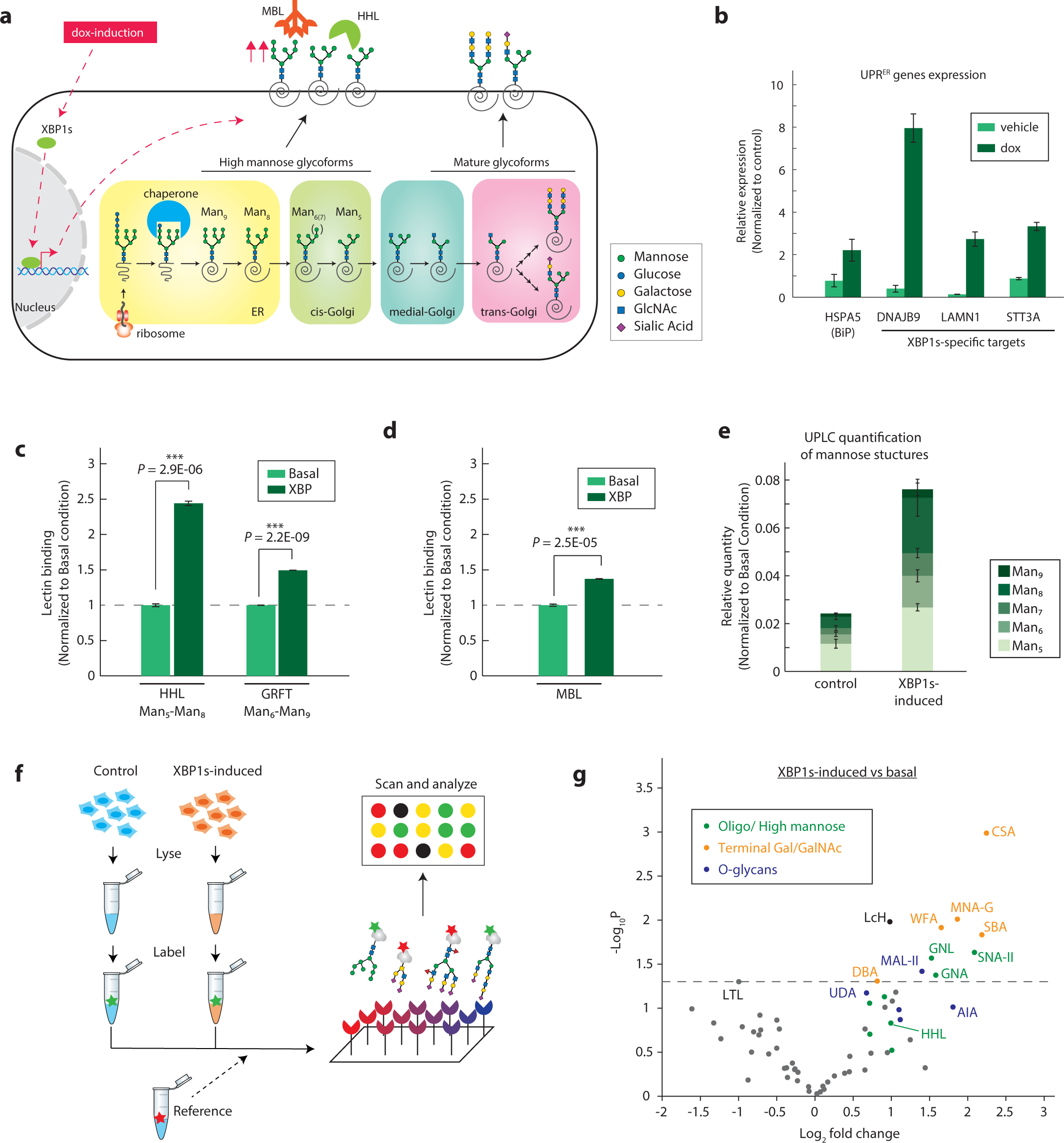
XBP1s-induction upregulates high mannose N-glycans on and within cells. **a,** Schematic for dox-inducible XBP1s upregulating high mannose N-glycans. **b,** RT-qPCR for targets of general UPRER and XBP1s. Gene expression is normalized to housekeeping genes GAPDH and HRPT1. Data are presented as mean ± s.e.m. and are representative of two independent experiments performed in triplicate with consistent results. **c,** Fluorescent HHL and GRFT binding on A549 cells with or without dox-induction of XBP1s. Cells were treated with 2 µg/mL dox for 48 hours to overexpress XBP1s. The flow cytometry data is representative of three independent experiments performed in triplicate. **d,** Fluorescent MBL2 binding on A549 cells with or without dox-induction of XBP1s. Cells were treated with 2 µg/mL dox for 48 hours to overexpress XBP1s. The flow cytometry data is representative of three independent experiments performed in triplicate. **e,** UPLC quantification of high mannose N-glycan structures of A549 cells with or without XBP1s-induction. Levels of each high mannose structure are normalized to the protein amount of each replicate. The experiment was performed in triplicate, and data is presented as mean ± s.e.m. **f,** Schematic for lectin microarray analysis of A549s under basal or XBP1s-induced conditions. **g,** Volcano plot of lectin microarray data. Median normalized log2 ratios (sample /reference) of the A549 samples are presented. Lectins are color-coded by their glycan-binding specificities.

Next, we tested whether XBP1s-induction by itself could alter cell surface high mannose glycan levels in A549. We utilized two lectins that can specifically bind high mannose N-glycan – Hippeastrum hybrid lectin (HHL), which binds N-glycan structures with Man_5_ to Man_8_^24^, and Griffithsin (GRFT), which binds Man_6_ to Man_9_^25^. We find that XBP1s activation leads to a slight but highly reproducible increase in both HHL and GRFT binding (**Fig. 1c, Extended Data Fig. 1b** and c). This increase in binding is reduced by cleaving off high mannose and hybrid glycans using Endoglycosidase H^3^, confirming the specificity of the lectins (**Extended Data Fig. 1d**). Notably, this increase in high mannose glycans on the cell surface also upregulates the binding of the complement pathway protein Mannose-binding lectin 2 (MBL2), consistent with previous reports^7^ (**Fig. 1c, Extended Data Fig. 1d**). In addition, partial activation of XBP1s using small molecule IXA4^26^ on wild type A549 cells also increases cell surface high mannose structures in a dose-dependent manner (**Extended Data Fig. 1f**). Together, these results show that activating the XBP1s branch of UPR^ER^ in A549 cells can enhance cell surface high mannose levels.

Next, to determine how each high mannose glycan structure (Man_5_ – Man_9_) changes upon XBP1s-induction on a whole cell level, we quantified all high mannose N-glycan structures using Ultra-Performance Liquid Chromatography (UPLC) with fluorescence detection^27^. Interestingly, XBP1s-induction massively upregulates all high mannose structures on a whole cell level, with the largest increase in Man 6-8 structures (**Fig. 1d, Extended Data Fig. 1g**). This data confirms that the high mannose expression in lung cells can be triggered by XBP1-pathway induction, providing a mechanism for glycan-based reporting of cell damage and infection to the innate immune system. Our results also suggest that the changes in cell surface glycome likely originate from changes in the early stages of N-glycosylation that occurs in the ER and Golgi.

Finally, to fully characterize the other glycosylation changes induced by XBP1s activation, we employed lectin microarray to comprehensively profile changes in glycan repertoire under basal and XBP1s-induced conditions (**Fig.1e, Source Data 1**). We confirmed the changes in high mannose structures (HHL) and observed corresponding upregulation of oligomannose structures (Man3 to Man9) by the increased binding of lectins SNA-II, UDA, and GNA^24^ (**Fig. 1f, Extended Data Fig. 1h**). Interestingly, we did not observe a corresponding decrease in complex glycans but instead uncovered a concurrent upregulation of complex N-glycans capped with terminal Galactose (Gal) and N-Acetylgalactosamine (GalNAc), suggesting that cells can upregulate high mannose independently as a stress signal without compromising complex N-glycan synthesis. This is consistent with findings in influenza infection, in which high mannose upregulation did not impact the expression of most complex glycan epitopes ^7^. Furthermore, we observe an upregulation of O-linked glycans (lectins: AIA, MNA, and MPL), highlighting how XBP1s induction globally alters cellular glycosylation.

### Genome-wide CRISPR screen uncovers the expanded network of genes regulating high mannose

To uncover the genes beyond glycan biosynthetic enzymes controlling cell surface high mannose levels, we utilized our cellular system to conduct a genome-wide CRISPR screen (Fig. 2a). To do so, we first engineered the XBP1s-inducible A549 line to also stably express Cas9 and confirmed that concurrent expression of sgRNA targeting relevant genes, such as XBP1, can alter cell surface presentation of high mannose glycans (**Extended Data Fig. 2A**).

**Fig. 2:**
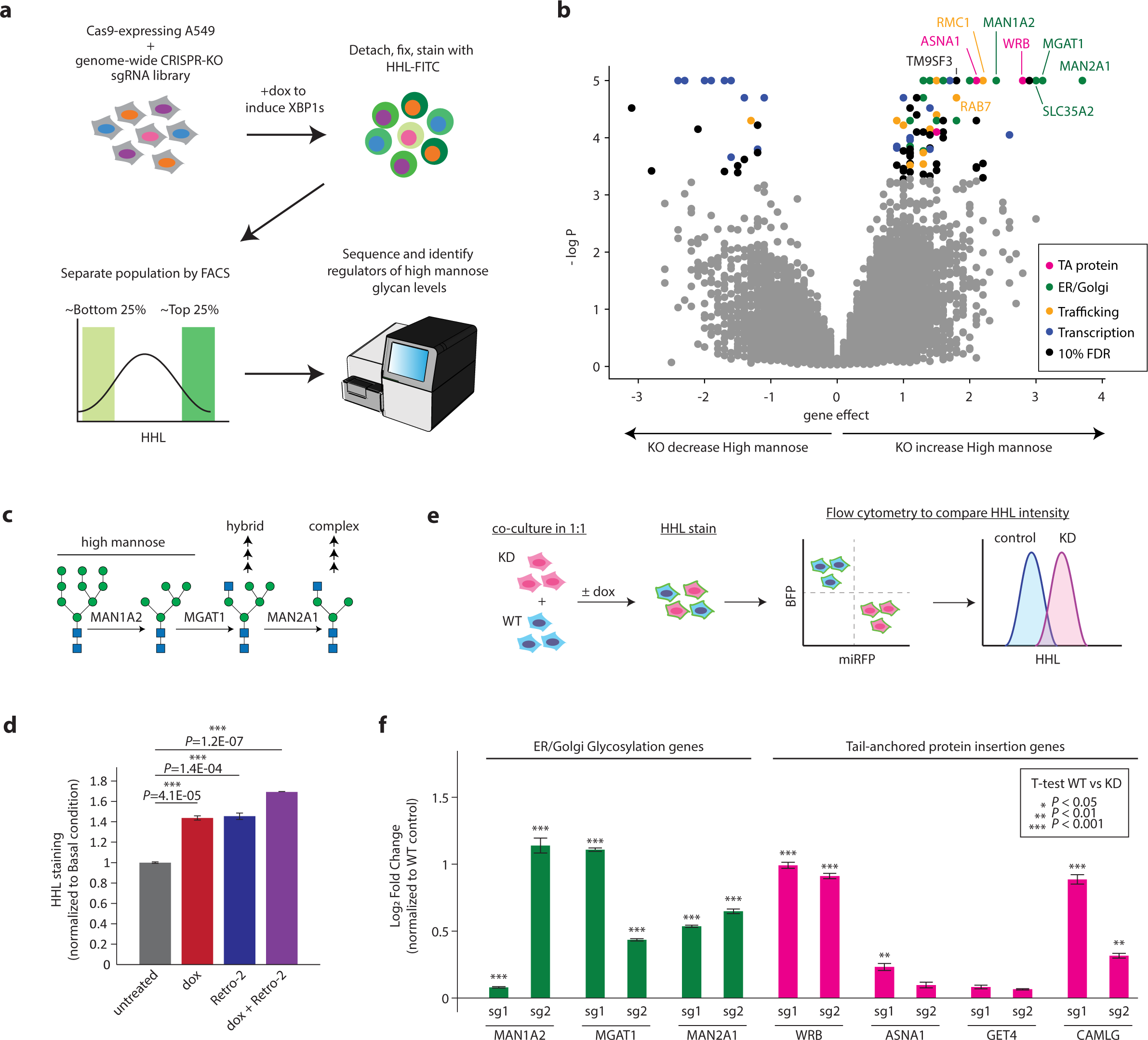
Genome-wide CRISPR screen uncovers the expanded network of genes regulating high mannose. **a,** Schematic for FACS-based CRISPR screen. Cas9-expressing A549s were lentivirally transduced with a genome-wide CRISPR-deletion sgRNA library. Resulting cells were dox-treated to induce XBP1s overexpression for 48 hours. Cells were then gently lifted with Accutase, fixed, and stained with FITC-labeled HHL. The top and bottom 25% of HHL stained cells were isolated by FACS. The resulting populations were subjected to deep sequencing and analysis. The screen was performed in duplicate. **b,** Volcano plot of all genes indicating effect and confidence scores for the genome-wide screen performed in duplicate. Effect and P values were calculated by casTLE. **c,** Schematic for initial steps of N-glycan mannose-trimming and remodeling. All three enzymes indicated are hits in genome-wide screen. **d,** Disruption of tail-anchored protein insertion pathway by ASNA1 inhibitor Retro-2 in wild type A549s also upregulates cell surface high mannose glycan levels. A549s were treated with treated with 2 µg/mL dox, 100 µM of Retro-2, both, or left untreated for 48 hours. Resulting cells were lifted with Accutase and stained with FITC-labeled HHL, followed by flow cytometry analysis. Data are presented as mean ± s.e.m. of median of each replicate and are representative of two independent experiments performed in triplicate with consistent results. **e,** Schematic for competitive binding assays for measuring changes in high mannose levels. Cells expressing sgRNAs for CRISPRi-mediated knockdown (KD) and miRFP and cells expressing a control sgRNA and BFP were cocultured in 1:1 ratio. Cells were either treated with dox to induce XBP1s or left untreated for 48 hours. Resulting cells were lifted and stained with HHL-FITC, and log2 ratio of HHL intensity of KO: control was determined using flow cytometry. **f,** Validation of hits in XBP1s-induced A549s using competitive HHL binding assays: Data are presented as mean ± s.e.m. and are representative of two independent experiments performed in triplicate with consistent results.

Next, we lentivirally transduced a previously validated genome-wide sgRNA knockout library^28^ into the Cas9-expressing, XBP1s-inducible A549 cells, with sgRNAs targeting all protein-coding genes with ten sgRNAs per gene and ∼10,000 negative controls. The cells were then dox-treated to induce XBP1s for 48 hours, fixed, and stained with FITC-labeled HHL. The population of cells with the top 25% and bottom 25% of HHL signal were selected using fluorescence-activated cell sorting (FACS), such that cells expressing sgRNAs targeting genes that suppress high mannose glycan presentation will be enriched in the top 25% and depleted in the bottom 25%, while cells expressing sgRNAs targeting genes required for high mannose glycans will be enriched in the bottom 25% and depleted in the top 25% population. The proportion of each sgRNA in the two populations was measured by deep sequencing, and significant regulators of high mannose glycan presentation were identified using casTLE^29^ (**Fig. 2a**).

This initial screen identified 109 known and novel regulators of high mannose glycan regulation at a 10% false discovery rate (**Fig. 2b, Source Data 2**). Among the strongest hits were biosynthetic enzymes directly involved in N-glycan maturation – MAN1A2 and MGAT1^22^. These are Golgi-localized enzymes that act in sequential steps to remove mannoses from Man_8_ and Man_9_ structures to form Man_5_ and add a GlcNAc residue (**Fig. 2c**). Deletion of any of these enzymes block glycan processing to more complex structures and can therefore lead to an increase in high mannose structures. Detection of these positive controls indicates that our screening strategy worked well to identify modulators of the high mannose epitope.

Besides enzymes involved in glycosylation, our strongest hits that enhanced high mannose glycan expression were members of the tail-anchored (TA) protein insertion pathway(**Fig. 2b**). Knocking out four of the six canonical members (WRB, GET4, ASNA1, and CAMLG) leads to an upregulation of high mannose structures. This is likely because disruption to the TA-insertion pathway mis-localizes essential Golgi proteins^30,31^, inhibiting proper N-glycan processing in the Golgi. Indeed, when we inhibited ASNA1 using the small molecule Retro-2^31,32^, cell surface high mannose glycans became upregulated under both basal and XBP1s-induced conditions (**Fig. 2d**). In contrast, many of the strongest genes that caused loss of high mannose upon deletion, even in the presence of induced XBP1s, were transcriptional regulators, some of which are likely to be involved in dox-induced overexpression of XBP1s and may not be involved in direct induction of high mannose glycans by XBP1s. Given this, we decided to focus on genes whose deletion enhanced high mannose levels in our assay regardless of XBP1s.

Next, we tested whether our top hits have the same impact on high mannose glycans under basal conditions. We established individual CRISPRi-knockdown cell lines with two independent sgRNA each and assayed how the disruption of each gene impacted cell surface high mannose levels using competitive HHL binding assays (**Fig. 2e**). We found that all our top hits regulate high mannose levels in both basal and XBP1s-induced conditions (**Fig. 2f, Extended Data Fig. 2c**). Together, these results highlight the critical role of Golgi function in regulating cell surface presentation of high mannose glycans.

### Magnetic sorting-based CRISPR screens uncover additional novel regulators of high mannose glycans under basal and UPR^ER^-induced conditions

We decided to focus our attention towards understanding high mannose regulation under basal, unstressed conditions. To do so, we generated a CRISPRi sublibrary targeting all genome-wide hits and genes functionally connected to our top hits, totaling 292 genes, with five sgRNAs each and 540 negative controls (Supplementary Table 1). Moreover, to enable faster screening at high coverage, we employed magnetic-activated cell sorting (MACS) to separate cells with high versus low levels of cell surface high mannose (**Fig. 3a**). We screened under both basal (untreated) or UPR^ER^ induced (XBP1s, dox) conditions to identify genes that impacted high mannose generally.

**Fig. 3:**
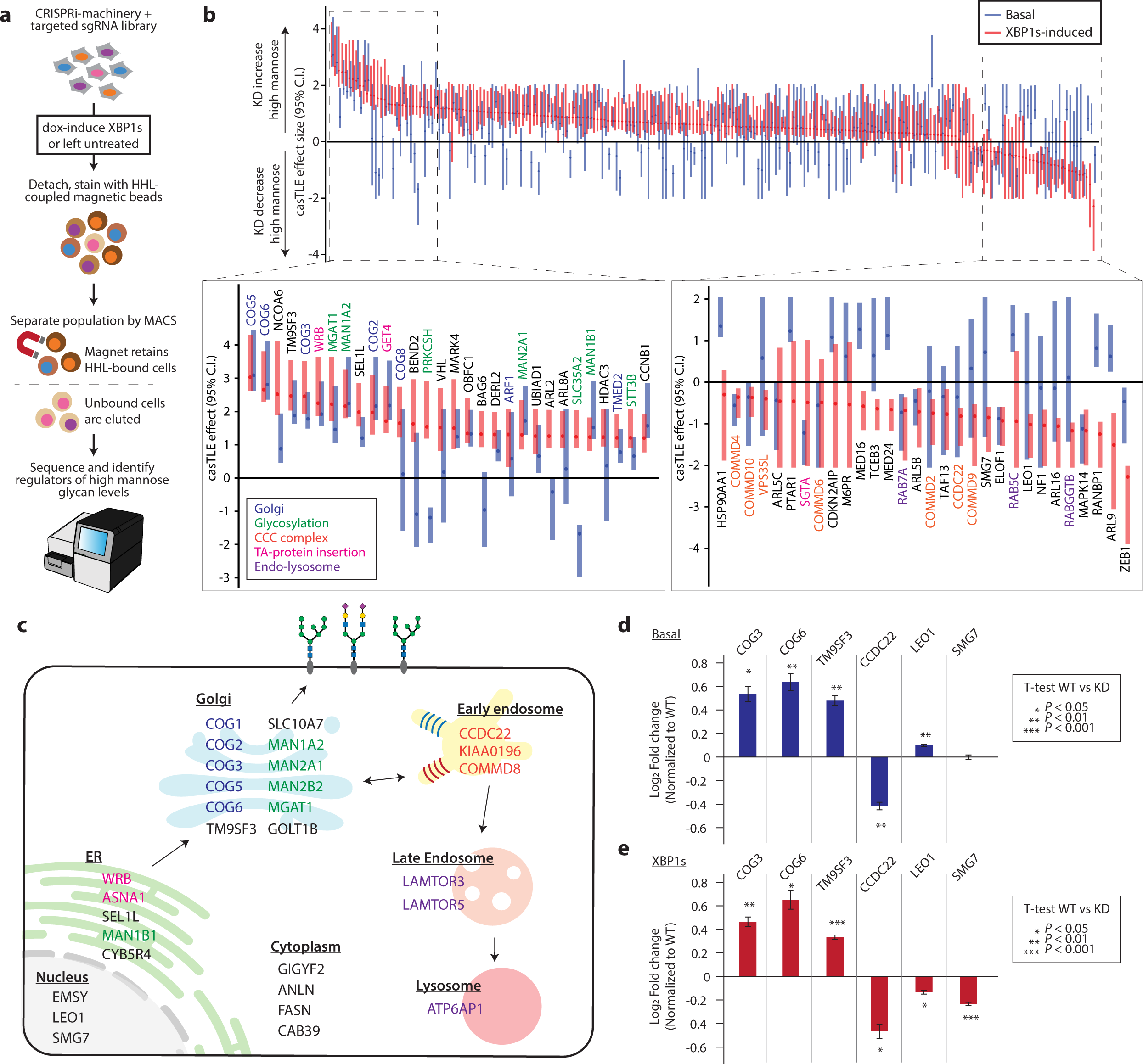
Targeted CRISPRi screen uncovers additional novel regulators of high mannose glycans under basal and UPR^ER^ induced conditions. **a,** Schematic for MACS-based CRISPR screen. A549 cells stably expressing CRISPRi machinery and the targeted sgRNA sublibrary were either dox-treated to induce XBP1s or left untreated for 48 hours. Cells were lifted and incubated with HHL coupled to magnetic beads. The cells were then placed on a magnet in which high HHL-binding cells would be retained on the magnet, whereas the low HHL-binding cells were removed from the population. This separation was repeated twice more on each high and low HHL binding cells to improve the purity of the populations. Finally, each resulting population were subjected to deep sequencing and analysis to identify hits. The screen was performed in duplicate. **b,** The maximum effect size (center value) estimated by CasTLE from both basal and XBP1s-induced conditions with five independent sgRNA per gene. The bars represent the 95% credible interval, with red representing XBP1s and blue representing basal conditions. Only genes considered to be a hit in at least one condition are shown. Genes are ordered in descending order of estimated maximum effect size of XBP1s-induced condition. The top 30 positive and negative hits are shown in the expanded panels. **c,** Top 30 regulators for high mannose N-glycans with their reported subcellular localization. **d,** Validation of hits in A549 under basal conditions using competitive HHL binding assays. Each gene is knocked down by co-expression of two independent sgRNAs. Data are presented as mean ± s.e.m. and are representative of two independent experiments performed in triplicate with consistent results. **e,** Validation of hits in A549 under XBP1s-induced conditions using competitive HHL binding assays. Each gene is knocked down by co-expression of two independent sgRNAs. Data are presented as mean ± s.e.m. and are representative of two independent experiments performed in triplicate with consistent results.

Briefly, the targeted sgRNA was lentivirally transduced into the A549 cell line with constitutively active CRISPRi machinery and dox-inducible XBP1s. The resulting cells were either treated with dox to induce XBP1s expression or left untreated. Cells were then lifted and stained with HHL conjugated to magnetic particles and separated magnetically such that cells with increased levels of high mannose on the cell surface would be retained by the magnet. In contrast, cells with less high mannose would be eluted. Each population was subjected to three rounds of separation. The proportion of each sgRNA was measured by deep sequencing and analyzed by casTLE^29^.

This strategy validated 77 hits from our genome-wide screen and further identified 111 additional genes that regulate the cell surface high mannose glycosylation under XBP1s induction. The increased number of genes identified in this secondary screen is likely due to the increased sensitivity from higher library coverage and CRISPR knockdown, which enabled essential genes to be more readily identified. Among these, 118 hits also regulate high mannose glycosylation under basal conditions. These include genes directly regulating early steps in processing the high mannose structure (e.g.MAN1A1, MAN1A2, and MGAT1). Identifying these glycosylation enzymes indicates that the screening approach worked well and has increased sensitivity in detecting high mannose regulators compared to the genome-wide screen. In addition, known Golgi regulators, such as all members of the COG complex (COG1-8), were also found to be regulators of the high mannose epitope (**Fig. 3b and c, Source Data 3**). Top hits were validated using competitive HHL binding assays (**Fig. 3d**).

Our strategy also enabled us to uncover genes that, when depleted, suppress high mannose levels (**Fig. 3b and c**). Interestingly, these include many members of the CCC protein complex, which consists of CCDC22, CCDC93, and any of the ten COMMD proteins. The CCC complex works closely with the retriever complex to regulate protein recycling between the endosome and the cell surface ^33,34^, but no glycosyltransferases have been reported as cargo. Given the tight connection between endocytic recycling and the trans-Golgi network^34^, it is plausible that the CCC might be acting through the Golgi to mediate high mannose levels. However, the precise role of the CCC complex in regulating high mannose or other types of glycosylation remains unclear.

As high mannose is a key intermediate for all N-glycans, we expected that many of our hits would impact other glycan structures along the N-glycan maturation pathway. Therefore, to evaluate how the top hits effect other forms of glycosylation, we measured changes in other glycan epitopes using a panel of lectins with known specificities^24^ on live, intact cells (**Extended Data Fig. 3c**). We found that knocking down known Golgi regulators (COG3, COG6, and GET1) generally shifts cells to display more high and oligomannose structures and fewer branched and complex epitopes. In contrast, disrupting the CCC complex (CCDC22 and VPS35L) leads to an upregulation of mature terminal glycan epitopes such as sialic acids and GalNAc and a corresponding downregulation of high and oligomannose structures. These results show that the top hits are not merely affecting glycan density on the cell surface but impacting the cell’s glycosylation pathways.

Together, our two-tiered screening approach allowed us to uncover known and novel regulators of high mannose glycosylation beyond expected biosynthetic enzymes under both basal and XBP1s-induced conditions. However, identifying the genes alone does not provide sufficient information for understanding how these regulators, particularly ones without known connection to the biosynthetic enzymes, control glycosylation. Thus, we next sought to investigate how two regulators of opposing phenotypes – TM9SF3 and the CCC complex – act to regulate glycosylation.

### TM9SF3 regulates Golgi organization and promotes N-glycan maturation

One of the strongest hits in our screens was a poorly characterized gene TM9SF3 (Transmembrane 9 Superfamily Member 3). Knock down of this geneleads to a strong upregulation of cell surface high mannose under both basal and XBP1s-induced conditions (**Fig. 3b, d, and e**). TM9SF3 belongs to a family of four multi-pass membrane proteins characterized by nine transmembrane domains and was previously found to be localized to the Golgi ^35,36^. Interestingly, its knockdown leads to similar glycan changes as known Golgi regulators (**Extended Data Fig. 3c**), suggesting that TM9SF3’s role may be linked to Golgi function. Moreover, another family member, TM9SF2, has been shown to regulate glycolipid synthesis^19,20^. However, the role of any TM9SFs in N-linked glycan regulation is unknown.

To begin understanding the mechanism by which TM9SF3 regulates high mannose glycosylation on the cell surface, we first validated its effect on high mannose by establishing three knockdown lines using independent sgRNAs and found that, as expected, all three lines have increased high mannose under both basal and XBP1s conditions (**Extended Data Fig. 4a**, b). We next we tested whether this was a general property of members of the TM9SF family. In line with our screen results, we find that only TM9SF3 acts to control high mannose levels on the cell surface (**Fig. 4a, Extended Data Fig. 4c**).

**Fig. 4:**
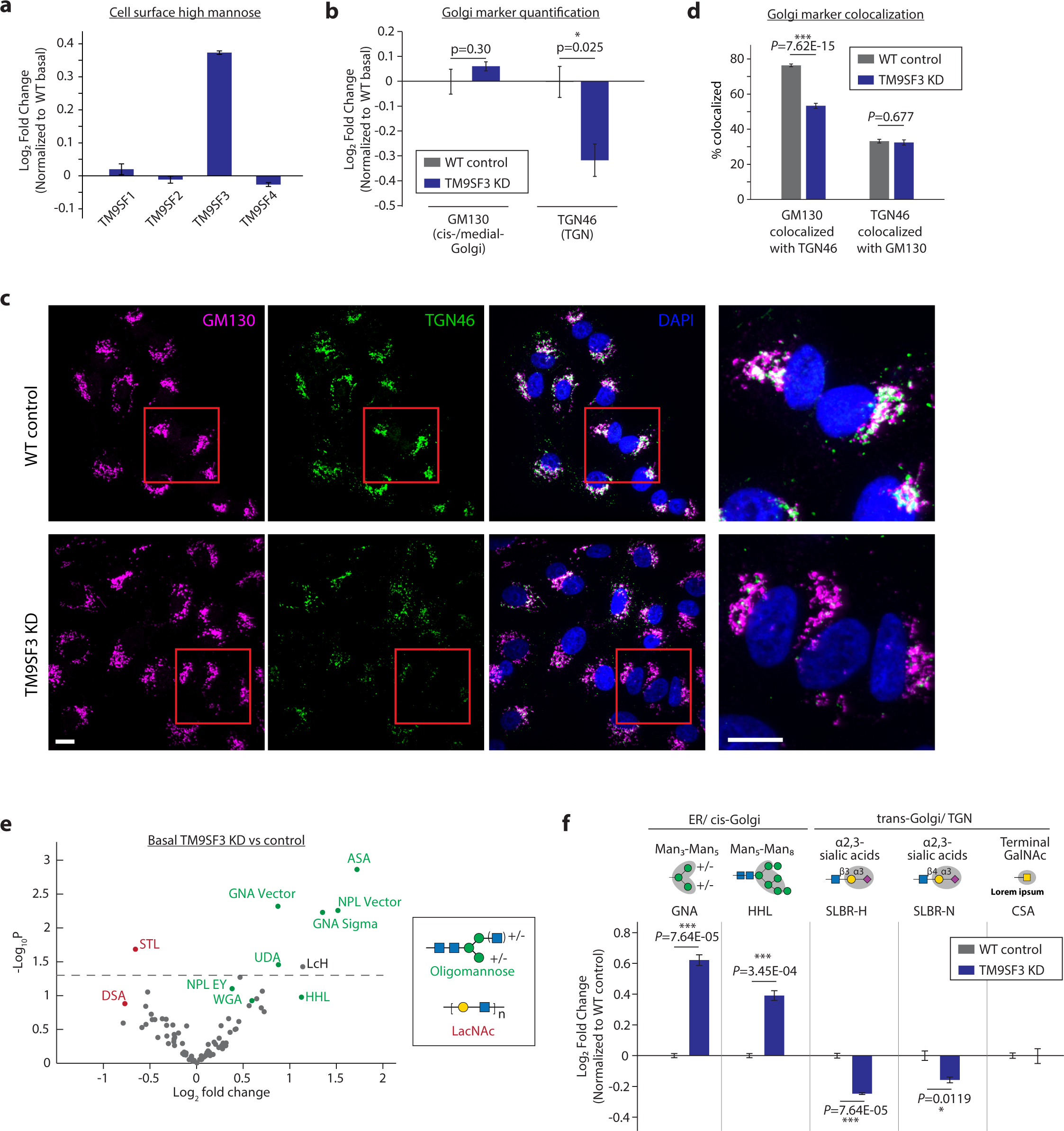
TM9SF3 regulates the Golgi organization and is required for formation of complex N-glycans. **a,** Competitive HHL binding assay in A549s with each TM9SF family member knocked down. Data are presented as mean ± s.e.m. and are representative of two independent experiments performed in triplicate. **b,** Flow cytometry quantification of intracellular staining of GM130 and TGN46 in A549s. Data are presented as mean ± s.e.m. and are representative of three independent experiments performed in triplicate. **c,** Representative confocal microscopy images of TM9SF3 knockdown and wildtype control cells, co-stained with cis-/medial-Golgi marker GM130 and TGN marker TGN46. Magnified views of the red boxed areas are shown in the right-most column. Scale bars, 10 µm. Images are representative of two independent experiments performed in triplicate. **d,** Percent area of each Golgi compartment co-localized with the other compartment. Data are presented as mean ± s.e.m., from 12 images each from wildtype or TM9SF3 knockdown of two independent experiments, with >20 cells per image. **e,** Volcano plot for lectin microarray results of A549 cells with TM9SF3 knocked down compared to wild type control. Lectins are color-coded by their glycan-binding specificities. **f,** Competitive cell surface lectin binding assay for TM9SF3 knocked down A549s compared to wild type control under basal conditions. Lectin binding specificities and location of where the modification predominately occurs are indicated. Data are presented as mean ± s.e.m. and are representative of three independent experiments performed in triplicate.

We reasoned that TM9SF3 might play a role in Golgi function and morphology. The Golgi has three compartments, the cis golgi-where mannosidases reside, the medial golgi, and the trans golgi network (TGN) where complex N-glycans and sialosides are synthesized. Using intracellular staining coupled with flow cytometry quantification to study TM9SF3 knocked down cells (TM9SF3-KD), we observe a slight decrease in TGN marker TGN46, suggesting that there might be mild defects in TGN function (**Fig. 4b, Extended Data 4d**). Characterization using confocal microscopy shows similar reduction in TGN46 staining (Fig. 4c). Interestingly, imaging also revealed changes in cis-and medial-Golgi morphology in TM9SF3-KD cells. Cis- and medial-Golgi compartments become highly dispersed, whereas TGN morphology and dispersion remains largely unchanged in TM9SF3-KD cells (**Fig. 4c and Extended Data Fig. 4f-h**). This leads to a reduction in cis-Golgi compartments that colocalize with TGN when compared to control cells (**Fig. 4d**). These results indicate that TM9SF3 regulates Golgi organization, which is essential for proper glycan maturation.

Next, we used lectin microarrays to unbiasedly measure changes in global glycosylation. Because the enzymes involved in glycan processing and their Golgi localization are largely known, obtaining a comprehensive survey of the glycan repertoire can provide mechanistic insights into what glycosylation steps, and potentially Golgi compartments, are altered when our gene-of-interest is knocked down. Surprisingly, this revealed a general upregulation of oligomannose glycans, suggesting that the early steps of N-glycan remodeling that converts high mannose into oligomannose structures, which occurs in the cis- and medial-Golgi, can proceed normally despite the altered morphology (**Fig. 4e, Extended Data Fig. 4j, Source Data 4**). In addition, we also observed a reduction in complex LacNAc epitopes, indicating that the final steps of glycan elongation and capping needed for forming complex glycans are inhibited. Indeed, in addition to a reduction in complex LacNAc epitopes, cell surface lectin binding assays also confirm the downregulation of other complex glycan epitopes such as α2,3-sialic acids (**Fig. 4f, Extended Data Fig. 4k**). Together, these results suggest that the fragmented cis- and medial-Golgi and the reduction in TGN in TM9SF3 knockdown cells may impede the trafficking of glycoproteins through Golgi compartments for glycan remodeling, resulting in a glycan repertoire enriched in high and oligomannose structures (**Extended Data Fig. 5j**).

### The CCC complex negatively regulates Golgi function and complex glycan formation

Finally, we sought to study the role of the CCC complex in regulating N-glycosylation, given that multiple complex members are identified as regulators of the high mannose epitope. To first validate our screen results and carefully determine how each complex member impacts the high mannose epitope, we established individual knockdown lines of each member using CRISPRi with two sgRNAs each. Consistent with the screen results, we find that knocking down its core members (CCDC22 and CCDC93) and 7 of its 10 COMMD members reduces high mannose epitope on the cell surface under both basal and XBP1s-induced conditions (**Fig. 5a, Extended Data Fig. 5a**). Notably, knocking down VPS35L, a component that the CCC shares with the Retriever complex^33^, also down-regulates high mannose and leads to a similar glycan profile as CCDC22 knockdown (**Extended Data Fig. 3c**), suggesting that the CCC and Retriever complex may act together to regulate glycosylation.

**Fig. 5:**
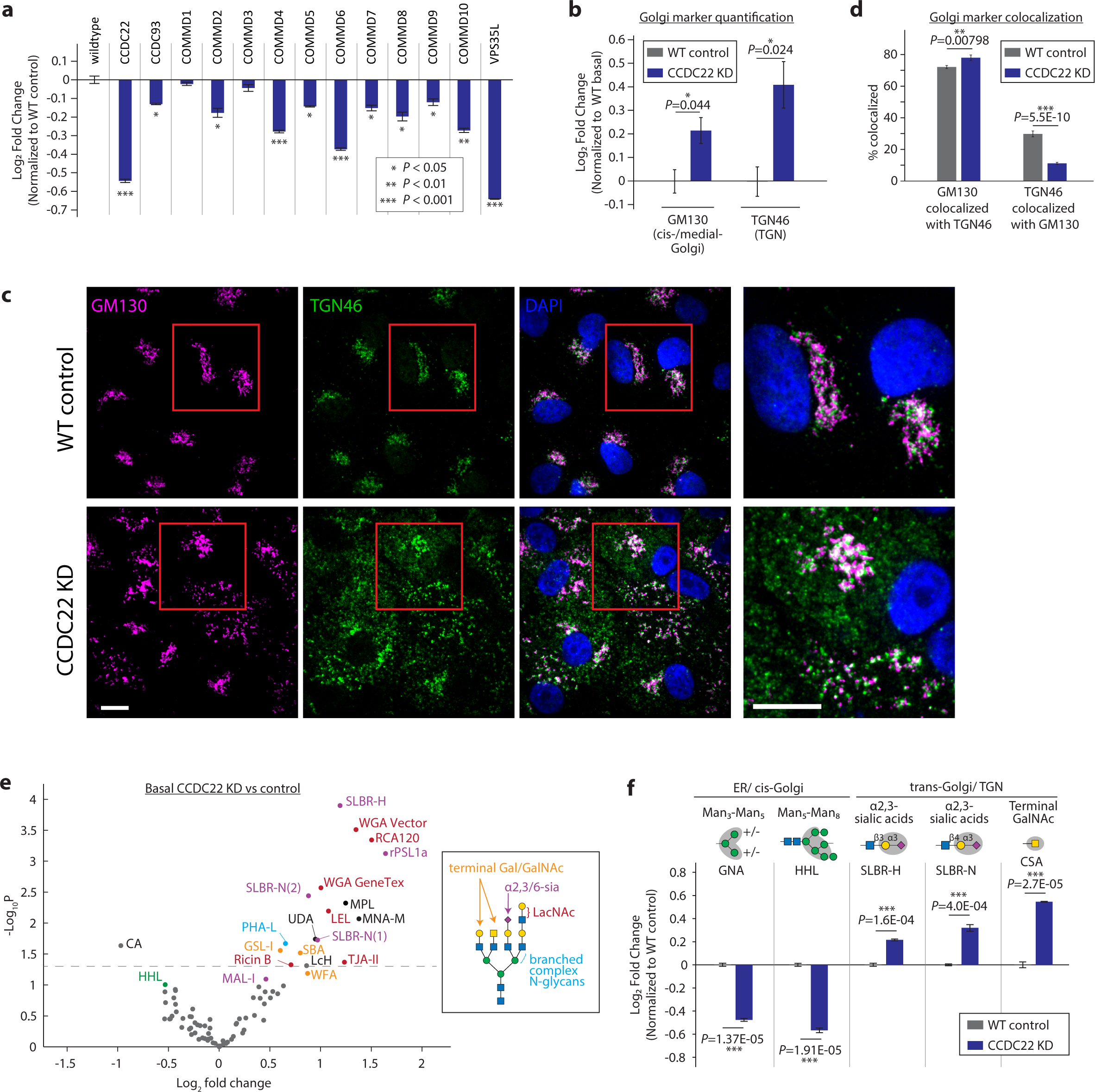
CCDC22 regulates elongation and sialylation of glycans through modulating Golgi expansion. **a,** Competitive HHL binding assays on A549s for all knock down of all members of the CCC complex. Each gene is knocked down by co-expression of two independent sgRNAs. Data are presented as mean ± s.e.m. and are representative of three independent experiments performed in triplicate. **b,** Flow cytometry quantification of intracellular staining of GM130 and TGN46 in A549s in CCDC22 knockdown and wild type control cells. Data are presented as mean ± s.e.m. and are representative of three independent experiments performed in triplicate with consistent results. **c,** Representative confocal microscopy images of CCDC22 knockdown and wildtype control cells, co-stained with cis-/medial-Golgi marker GM130 and TGN marker TGN46. Magnified views of the red boxed areas are shown in the right-most column. Scale bars, 10 µm. Images are representative of three independent experiments performed in triplicate. **d,** Percent area of each Golgi compartment co-localized with the other compartment. Data are presented as mean ± s.e.m., from 12 images each from wildtype or CCDC22 knockdown of two independent experiments, with >20 cells per image. **e,** Volcano plot for lectin microarray results of basal A549 cells with CCDC22 knocked down compared to wildtype control. Lectins are color-coded by their glycan-binding specificities. **f,** Competitive cell surface lectin binding assay for CCDC22 knocked down A549s compared to wildtype control under basal conditions. Lectin binding specificities and the location of where the modification predominately occurs are indicated. Data are presented as mean ± s.e.m. and are representative of two independent experiments performed in triplicate.

Given the critical role of protein recycling in the secretory pathway, we next sought to test how the Golgi might be impacted by disruption to the CCC complex. To do so, we generated a stable A549 line with essential CCC complex component CCDC22 knocked down (**Extended Data Fig. 5b**). We found that CCDC22-depletion leads to a slight upregulation in both cis-/medial-Golgi marker GM130 and TGN marker TGN46 (**Fig. 5b, Extended Data Fig. 5c**). To characterize these Golgi changes further, we used confocal microscopy to monitor cis-/medial-Golgi as well as TGN morphology in CCDC22 knockdown cells. Surprisingly, we observed a dispersed cis/medial-Golgi phenotype, similar to that observed in TM9SF3-KD cells despite the opposing phenotypes (**Fig. 5c, Extended Data Fig. 5d** and e). In the CCDC22-KD cells, we also observed a similar fragmentation and dispersion of the TGN (**Fig. 5c, Extended Data Fig. 5f** and g), which was consistent with previous reports^37^. Interestingly, despite the dispersion of the Golgi compartments, the cis-Golgi becomes even more colocalized with the TGN (**Fig. 5c and d**). These findings led us to hypothesize that disrupting the CCC complex might either (1) trap high mannose glycans intracellularly, preventing them from reaching the cell surface, or (2) enhance high mannose remodeling into more complex glycans through increased association between the Golgi compartments.

To test these possibilities, we utilized lectin microarray to thoroughly interrogate glycosylation and Golgi function. This revealed a dramatic upregulation of complex glycan epitopes –Glycans of CCDC22 knockdown cells are more likely to be highly branched, elongated with N-acetyllactosamine (LacNAc), and capped with terminal sialic acids or galactose (**Fig. 5e, Extended Data Fig. 5h, Source Data 4**). These changes are matched by their cell surface staining (**Fig. 5f, Extended Data Fig. 5i**). These findings strongly suggest that disruption of the CCC complex enhances the process by which high mannose glycans are remodeled into more complex N-glycans, resulting in an upregulation of complex glycans at the expense of high mannose glycans (**Extended Data Fig. 5k**). This may be due to the increased association between the Golgi compartments, allowing glycoproteins to be more efficiently trafficked through the Golgi and thereby promoting glycan maturation. The expanded Golgi network may also be concentrating glycan synthesis enzymes, particularly elongation and capping glycosyltransferases to generate more complex glycans. Together, our results indicate that the CCC complex is a negative regulator of Golgi function and complex N-glycan formation.

## Discussion

In this study, we present an approach utilizing CRISPR screening and lectin microarrays to identify and characterize the network of genes that regulate cell surface glycosylation. Applying this strategy, we first used genome-wide and targeted CRISPR screens to uncover regulators of high mannose glycosylation, which enabled us to identify genes beyond the known biosynthetic enzymes. We then used lectin microarrays to comprehensively measure glycosylation changes in two novel regulators – a previously uncharacterized gene TM9SF3, and the protein recycling machinery CCC complex. Our analyses indicate that TM9SF3 is a regulator of Golgi organization and is required for proper complex N-glycan synthesis, whereas the CCC complex is revealed to be a negative regulator of Golgi function and complex glycosylation.

While it is no surprise that regulators of the Golgi would influence high mannose and other types of glycosylation, our approach allowed us to identify regulators of Golgi function in a manner traditional morphology or single glycoprotein analyses did not provide. Notably, the use of lectin microarrays allowed us to rapidly measure changes in N- and O-glycans simultaneously, providing comprehensive insights into the state of glycosylation pathways in the cell without the need to follow specific glycosyltransferases, which can be technically challenging due to their overlapping functions as well as low protein expression. Specifically, our work found an unexpected disconnection between Golgi morphology and function, in which fragmented and dispersed Golgi appear to retain function to process glycans. Interestingly, these scattered, smaller Golgi structures observed in our TM9SF3 and CCDC22 knockdown cells are reminiscent of Golgi satellites or outposts in dendrites of neurons, where localized glycosylation events can occur in response to neuronal excitation^26,38^, suggesting that such regulation of Golgi morphology and function may be a general mechanism by which cells control glycosylation. Specifically, fragmentation and dispersion only in the cis- and medial-Golgi along with the disconnection from the TGN, like what we observe in TM9SF3-KD cells, might allow high and oligomannose glycans to bypass the intact trans-Golgi deplete cells of their complex glycan structures. On the other hand, fragmentation and rearrangement of the Golgi to bring cis- and trans-golgi into closer contact, as we observed when the CCC complex is disrupted, might allow for concentrating specific glycosyltransferases and/or more efficient trafficking through the Golgi, enhancing remodeling and upregulating complex glycans. Follow-up studies will be required to determine whether these genes regulate glycome changes in disease states and fully elucidate how they may dynamically regulate glycosylation enzymes and glycosylation of specific proteins.

Glycosylation changes brought by changes in Golgi dynamics can have significant implications for how cells interact with the immune system. Particularly, the upregulation of complex epitopes capped with sialic acids has been shown to suppress the immune system^39^. In contrast, excess high mannose epitopes can over-activate complement pathways through interaction with mannose-binding lectin ^7,8^ and promote cancer metastasis ^40^. Notably, Golgi dysregulation is a feature of many diseases, including bacterial and viral pathogenic infections, cancers, and neurodegenerative diseases^41–43^. Careful research into how the Golgi alterations in various diseases regulate and change glycosylation can provide insight into how they may alter cell surface glycan signals to escape surveillance or overactivate the immune system to cause chronic inflammation.

Together, our work discovered novel regulators of high mannose glycosylation and Golgi function. Additionally, our work demonstrates a readily generalizable approach combining CRISPR screens and lectin microarrays for dissecting the complex network of genes that controls the production of any glycan epitopes. This can be easily adapted to different cell types for studying the cell-type specificity of glycosylation^12,44^. Collectively, this represents a powerful method for understanding glycosylation regulation and can allow us to investigate the origins of altered glycosylation in many diseases.

## Methods

### Cell culture

A549 and 293T cells were obtained from UC Berkeley Cell Culture Facility. A549s were grown in DMEM (Gibco 11966025) supplemented with 10% fetal bovine serum (FBS, Avantor, 97068-085), and 1% penicillin-streptomycin (Gibco, 15070063). 293Ts were grown in DMEM supplemented with 10% FBS, 2mM glutamax (Gibco 35050061), and 1% penicillin-streptomycin. All cells were cultured at 37°C with 5% CO_2_.

### RT-qPCR for UPR^ER^ targets

A549 cells with dox-inducible Cas9 were either treated with 2 µg/mL doxycycline (Sigma, D3072) or equal volume of DMSO for 48 hours. Cells were washed 2x with ice cold PBS (Gibco, 10010049). TRIzol reagent (Invitrogen, 15596026) and RNeasy micro kit (Qiagen catalog no. 74004) were used in conjunction to isolate cellular RNA according to manufacture instructions. cDNA synthesis from the purified RNA was performed using QuantiTect Reverse Transcription kit (Qiagen, catalog number 205311) according to manufacturer’s instructions. SYBR Green master mix (Applied Biosystem, catalog number A46109) and primers listed below were used to set up qPCR reactions and analyzed on QuantStudio 6 Flex.

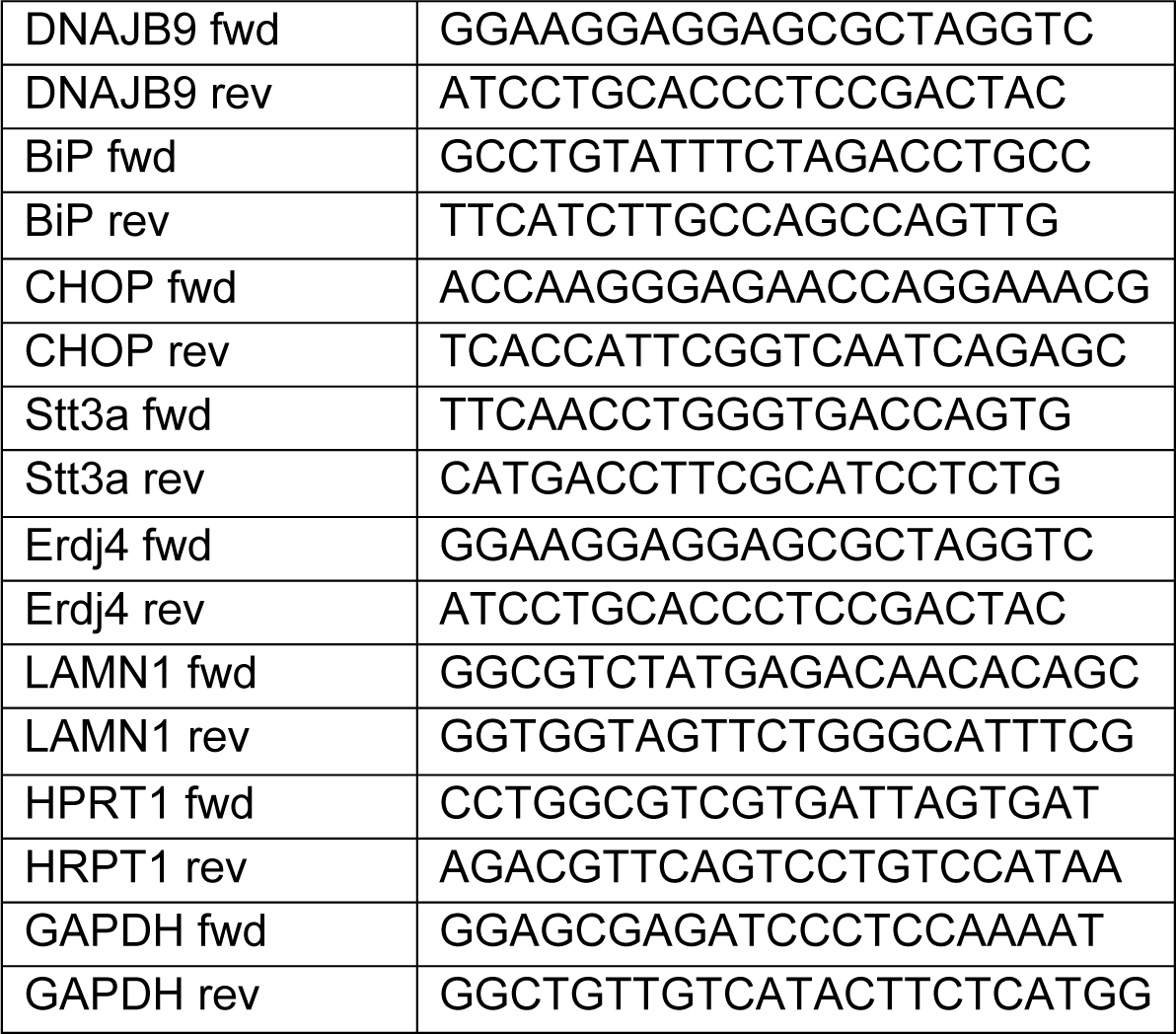

### Flow cytometry analysis of lectins and MBL2 binding

Cells were washed 2x with PBS and gently dissociated with 1:3 diluted Accutase (Gibco, A11105-01). Lifted cells were washed 2x with cold dPBS (HyClone, SH30264.02) and incubated with lectins at 10 µg/mL or MBL2 protein (Abcam, ab151947) at 1:50 in 3% bovine serum albumin (BSA) on ice for 1 hour. HHL was conjugated with FITC (Glycomatrix, catalog number 21511092-1), whereas His-tagged GRFT made in-house in the Mahal lab, and pre-incubated with Alexa fluor 488-labeled anti-His antibodies (Biotechne, IC0501G). After incubation with lectins or MBL2, cells were washed 2x with cold dPBS and analyzed on Attune NxT using the BL2 channel for green fluorescence. Other lectins used in flow cytometry experiments include: GNL (Vector Laboratories, FL-1241-2), Lch (EY Laboratories, F-1401-5), SNA (Bioworld 21500045-1), PNA (XXXX),CSA (EY Laboratories, BA-3201-1), PHA-L, SLBR-H, and SLBR-N (made in-house in the Mahal Lab).

### UPLC Quantification of high mannose N-glycan structures

Proteins were harvested from A549s with or without XBP1s-induction using 1% NP-40 lysis buffer -- 1% NP-40, 150 mM NaCl, and 50 mM Tris-Cl pH 8) supplemented with cOmplete protease inhibitor (Roche, 11836170001). Isolated proteins were flash frozen and sent to UC San Diego’s GlycoAnalytic Core for N-glycan analysis using Ultra-Performance Liquid Chromatography (UPLC) with fluorescent detection. Briefly, N-glycans were cleaved off by PNGase F, purified, and labeled with procainamide to allow for detection. The same amount of high mannose N-glycan structures with Man_5_-Man_9_ were spiked into each sample to allow for relative quantification of each high mannose N-glycan structure.

### Lectin microarrays

Flash frozen A549 cell pellets were washed with protease inhibitor cocktail supplemented PBS and sonicated on ice until homogenous. 20 µg of protein from each homogenized samples were then labeled with Alexa Fluor 555-NHS. A reference sample was prepared by pooling equal amounts (by total protein) of all samples and labeled with Alexa Fluor 647-NHS. Lectin microarray printing, hybridization, and data analysis was performed as previously described^14^. Details for the print are provided in the MIRAGE table (Supplementary table 4).

### FACS-based CRISPR-deletion screen for high mannose regulators

A previously established genome-wide, 10 sgRNA per gene CRISPR deletion library which is separated into 9 sublibraries was used for the genome-wide screen. A549s confirmed to stably express Cas9 and the dox-inducible XBP1s circuit was transduced with one sublibrary at a time at a multiplicity of infection (MOI) of 0.3-0.4. Cells expressing sgRNAs were selected using puromycin (Gibco, A1113803) at 1 µg/mL for 3-4 days such that >90% of cells were mcherry positive as measured by flow cytometry. Cells were then allowed to expand for up to 7 days. Deep sequencing was used to confirm sufficient sgRNA representation in each library.

The screen was performed one sublibrary at a time due to the large amount of FACS required. For each sublibrary, cells were treated with 2 µg/mL dox for 48 hours to induce XBP1s expression. Cells were dissociated with Accutase and fixed with 4% PFA. Fixed cells were stained with HHL-FITC at 10 µg/mL in 3% BSA for 2 hours at 4C with rotation. Cells were then washed 2x with dPBS and resuspended in 3% BSA. HHL-stained cells were sorted on BD Aria for top and bottom 25% of HHL signal, with at least 1000x coverage in each population. Cells were sorted within a week. The recovered cells were unfixed by incubating with protease K (Qiagen, 19133) overnight at 56°C with shaking. Genomic DNA of each population was extracted using Qiagen DNA Blood Midi kit (Qiagen, 51183). The sgRNAs were amplified and prepared for sequencing with a previously described nested PCR protocol with slight modification to make sgRNA sequencing library compatible with Illumina read 1 primer. Briefly, the sgRNA-encoding constructs were first amplified with primers oKT187 and oKT188, followed by a second PCR to introduce staggered sequences and indices for multiplexing (see Supplementary Table 2 for primer sequences). The resulting PCR products were gel purified prior to sequencing on Illumina HiSeq. Hit identification was performed using CasTLE^29^. See Supplementary Table 3 for all sgRNAs used for validation.

### MACS-based CRISPR-inhibition screens

To generated CRISPRi A549s, a CRISPRi construct (pLX_311-KRAB-dCas9, gift from John Doench & William Hahn & David Root, Addgene plasmid # 96918; http://n2t.net/addgene:96918; RRID:Addgene_96918) was lentivirally introduced into the A549s expressing inducible-XBP1s. These cells were selected with blasticidin (10 µg/mL) and single cell cloned to ensure stable CRISPRi machinery expression. To generate the secondary CRISPRi screening library, sgRNAs targeting a total of 292 genes, including all genes that passed 10% FDR from the genome-wide screen as well as functionally related genes were designed using CRISPick^45,46^, along with ∼500 control sgRNAs were synthesized by Twist Bioscience and cloned into pMCB320 using BstXI/BlpI overhangs after PCR amplification (see Supplementary Table 1 for complete list of genes and sgRNAs). This library was lentivirally installed into the A549s expressing CRISPRi machinery as well as inducible-XBP1s, and selected for with puromycin (1 µg/ mL).

For the screen, the 50 million library cells per condition were seeded in ten 15 cm plates. Cells were either treated with dox for 48 hours to induce XBP1s epxression or equal volume of DMSO as basal control. Cells were then lifted with Accutase, pooled, and washed 2x with dPBS. Cells were then resuspended in 10 mL 1% BSA and incubated with 100uL HHL-coupled with magnetic beads, which were prepared by mixing biotinylated HHL and MojoSort streptavidin nanobeads (BioLegend, 480016) 1:1 for 30 minutes on ice. HHL-beads were allowed to bind to cells for 1 hour at 4°C with rotation. Cells were then washed 2x with cold MojoSort buffer (BioLegend, 480017), and resuspended in MojoSort buffer. Cells were then placed on magnet and allowed to separate for 10 minutes. Unbound cells were collected into new tubes, whereas bound cells were resuspended in MojoSort buffer and allowed to separate again. After 10 minutes of separation, the unbound population is discarded, and the bound population was resuspended and allowed to separate one more time to increase purity. Similarly, the initial unbound population was placed on the magnet again, allowed to separate twice more by keeping the unbound population and discarding the bound population. Each bound and unbound populations underwent a total of three rounds of separations. A total of 10M and 20M cells were collected for the high mannose-low and -high populations, representing a 5000x and 10000x coverage of the sgRNA library, respectively Finally, genomic DNA were extracted from all resulting populations using Qiagen Blood Midi Prep and sgRNAs were prepared for sequencing in the same manner as the genome-wide screen. Hit identification were performed using CasTLE^29^. See Supplementary Table 3 for all sgRNAs used for validation.

### Intracellular staining and flow cytometry quantification of GM130 and TGN46

Cells were stained with standard intracellular staining techniques. Briefly, cells gently dissociated with 1:3 diluted Accutase, washed, fixed with 4% PFA (4°C, 15 minutes), permeabilized with 0.1% Triton X-100 (room temperature, 15 minutes), and blocked with 5% FBS in dPBS (room temp, 1 hour). They were stained for 2 hours at room temp with the following primary antibodies: mouse anti-GM130 (1:250, BD Bioscience, BDB610822) and rabbit anti-TGN46 (1:500, ProteinTech, 13573-1-AP), followed by 30 minutes of incubation with the following secondary antibodies: Goat anti-mouse (1:1000, Invitrogen, A-11015), Goat anti-rabbit (1:1000, Invitrogen, A-11011). The fluorescence signal was quantified on Attune NxT.

### Immunofluorescence and confocal microscopy

Cells were grown on glass coverslips were stained using standard immunocytochemistry techniques. Briefly, cells were fixed with 4% PFA, permeabilized with 0.1% Triton X-100, blocked with 3% BSA and stained with the following antibodies: mouse anti-GM130 (1:250), rabbit anti-TGN46 (1:500), and Phalloidin Alexa Fluor 647 (Thermofisher, A30107). Cover slips were mounted using VectaShield with DAPI (Vector Laboratories, H-1800-2). All images were collected on a Nikon Ti-E inverted microscope (Nikon Instruments, Melville, NY) equipped with a Plan Apo 60× oil objective. Images were acquired using a Zyla 5.5 camera (Andor Technology), using the iQ3 acquisition software (Andor Technology).

## Data Availability

The complete results of genome-wide screens and secondary screens are in Source Data related to Fig 2 and 3. The sgRNA counts of the screens are available through Figshare. The complete results of lectin microarrays are in Source Data related to Fig. 1 and 4. All data are available from the corresponding author upon reasonable request.

## Code Availability

casTLE v.1.0 is available at https://bitbucket.org/dmorgens/castle.

## Supporting information

Source Data for Fig. 1f

Source Data for Fig. 2b

Source Data for Fig. 3b

Source Data for Fig. 4e and 5e

## Acknowledgments

We thank members of the Dillin lab and D. Morgens for feedback on the project and manuscript; members of the UC Berkeley Flow Cytometry Core for assisting in FACS; members of the UC San Diego GlycoAnalytics Core for performing the high mannose N-glycan analysis; members of the UC Berkeley QB3 Sequencing Core for sequencing the genome-wide screen libraries; L. Hanqin and G. Pangilinan of the laboratory of Dirk Hockemeyer for sequencing of the targeted library screens; A. Jain and R. Zoncu for assistance on confocal microscopy experiments. This work was funded by NIH (grant no. 5F32AG069388-03 to C.K.T.), Howard Hughes Medical Institute (A.D.), and the Canada Excellence Research Chair Program (L.K.M.). We also acknowledge that this work used services provided by the GlycoNet Integrated Services.

## Author information

Authors and Affiliations

**Department of Molecular and Cell Biology, Howard Hughes Medical Institute, University of California, Berkeley, Berkeley, CA 94720, USA**

C. Kimberly Tsui, Emma Doan, Jacqueline Woo, Noosha Khosrojerdi, Janiya Brooks, Brant Webster, Andrew Dillin

**Department of Chemistry, University of Alberta, Edmonton, Canada, T6G 2G2**

Nicholas Twells, Ayodeji Kulepa, Lara K Mahal

## Contributions

C.K.T. conceived the project, designed, performed the majority of the experiments, analyzed data, and wrote the manuscript. N.T. performed the lectin microarray experiments. N.T. and A.K. generated recombinant lectins (GRFT, SLBR-H, and SLBR-N). E.D. J.W., N.K, and J.B. assisted in sgRNA validations. B.W. provided cloning plasmids for XBP1s constructs. A.D. and L.K.M. supervised the study.

## Supplementary Information

**Supplementary Tables 1**

Supplementary Table 1: Complete list of genes and sgRNAs for MACS-based screen. Supplementary Table 2: Primers used for CRISPR screens library prep. Supplementary Table 3: All sgRNAs used for validation experiments.

**Supplementary Table 4**

MIRAGE table for lectin microarray print details.

---

**Extended Data Fig. 1.**
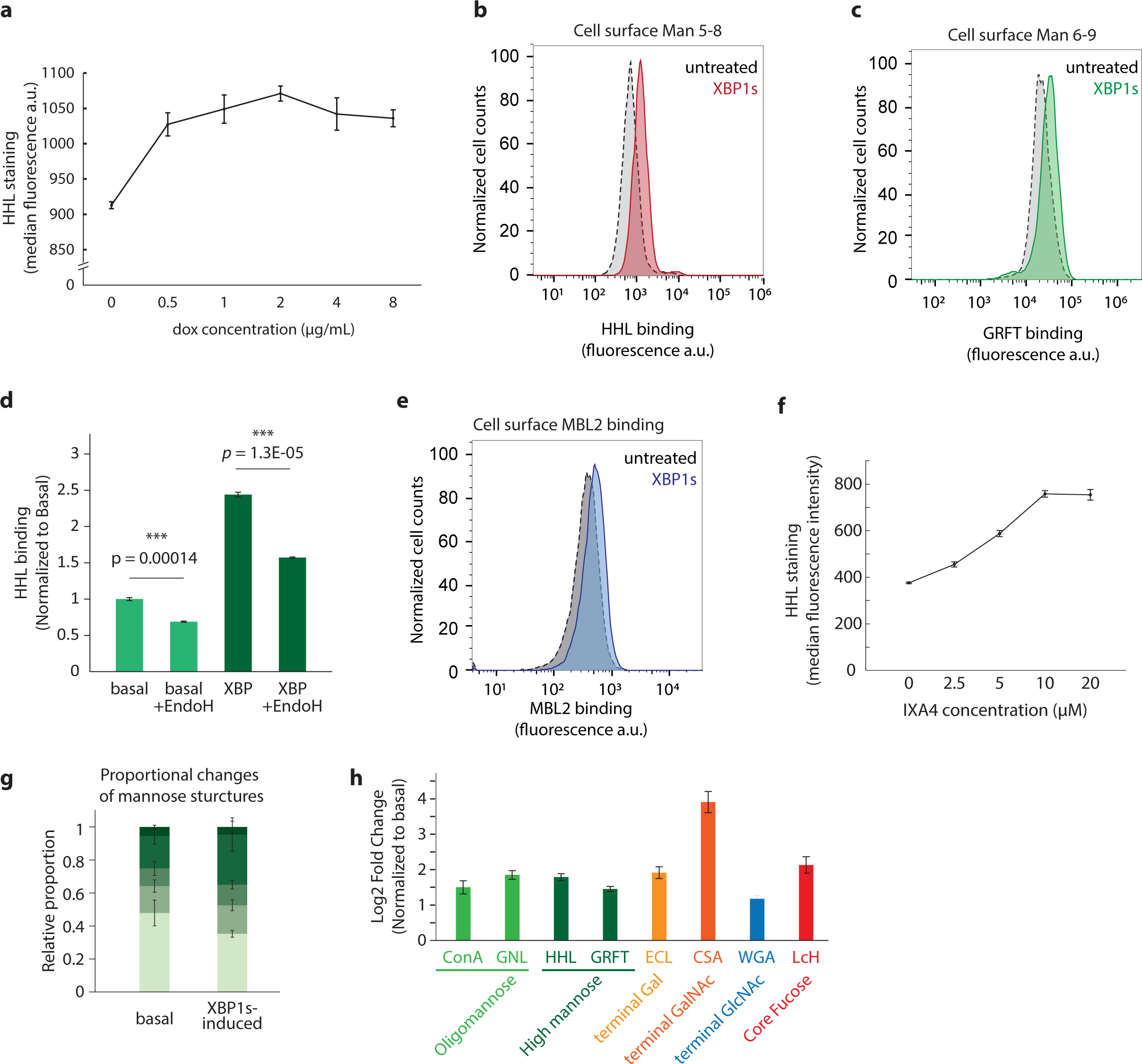
**a,** Dox-titration of XBP1s-induction. Cell surface HHL-binding is measured by flow cytometry. Data are presented as mean ± s.e.m. and are representative of two independent experiments performed in triplicate with consistent results. **b,** Representative flow cytometry results of lectin binding for HHL as related to Fig. 1c. **c,** Representative flow cytometry results of lectin binding for GRFT as related to Fig. 1C. **d,** HHL staining on cells treated with Endoglycosidase H for 1 hour at 37°C. Data are presented as mean ± s.e.m. and are representative of two independent experiments performed in triplicate with consistent results. **e,** Representative flow cytometry results of lectin binding for MBL2 as related to Fig. 1C. **f,** Cell surface HHL-binding on cells treated with increasing concentration of IXA4. Flow cytometry results are presented as mean ± s.e.m. and are representative of two independent experiments performed in triplicate with consistent results. **g,** Proportional changes of high mannose structures in A549 cells with or without XBP1s-induction, as related to Fig. 1d. The experiment was performed in triplicate, and data is presented as mean ± s.e.m. **h,** Cell surface lectin binding of XBP1s-induced cells normalized to basal control cells. Data are representative of two independent experiments performed in triplicate.

**Extended Data Fig. 2.**
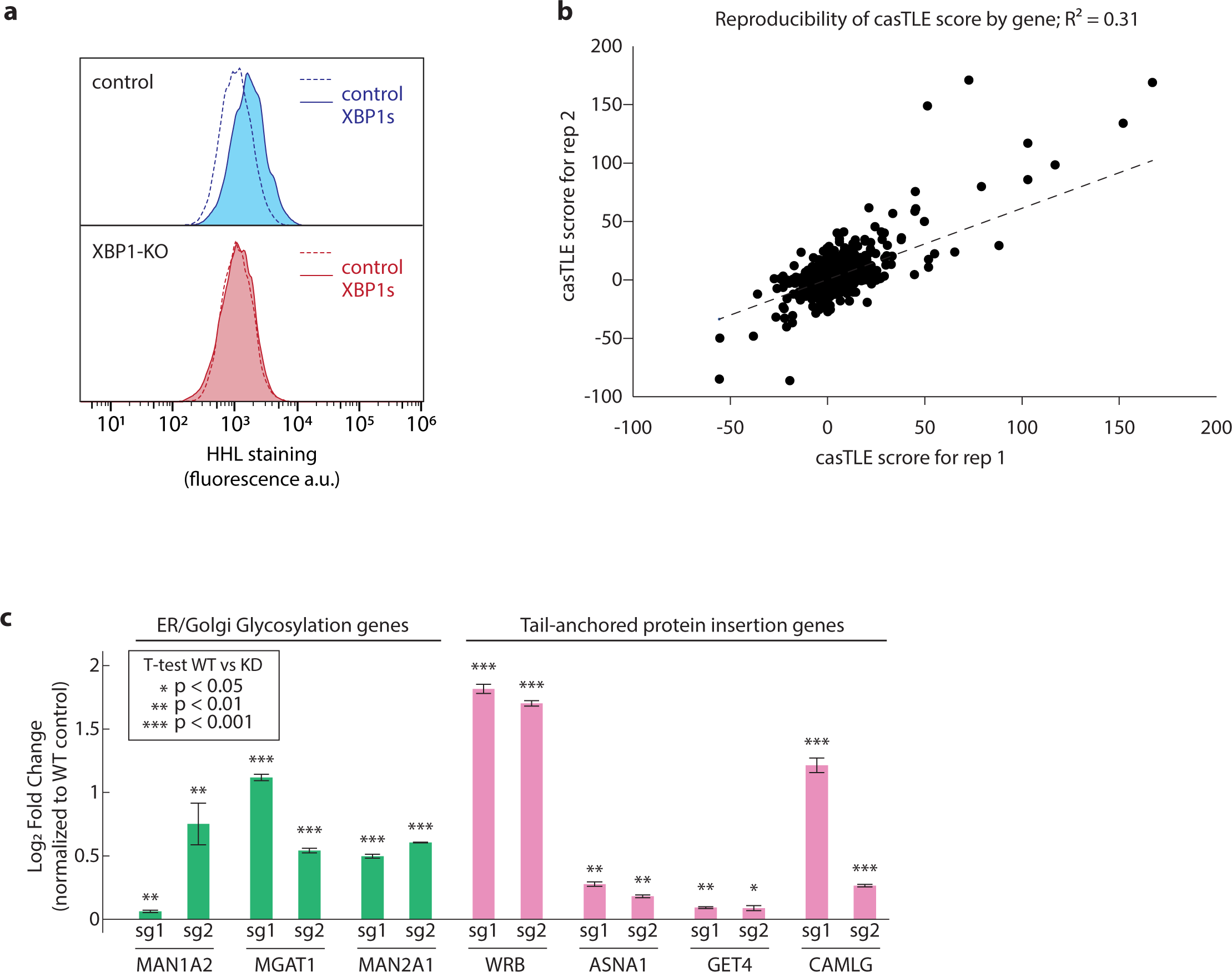
**a,** Flow cytometry result of HHL binding in A549 wild type control and cells expressing XBP1-targeting sgRNA (XBP1-KO). Data are representative of two independent experiments performed in triplicate. **b,** Reproducibility plot of casTLE scores for the CRISPR-KO genome-wide screen. **c,** Validation of hits in basal A549s using competitive HHL binding assays: Data are presented as mean ± s.e.m. and are representative of two independent experiments performed in triplicate with consistent results.

**Extended Data Fig. 3.**
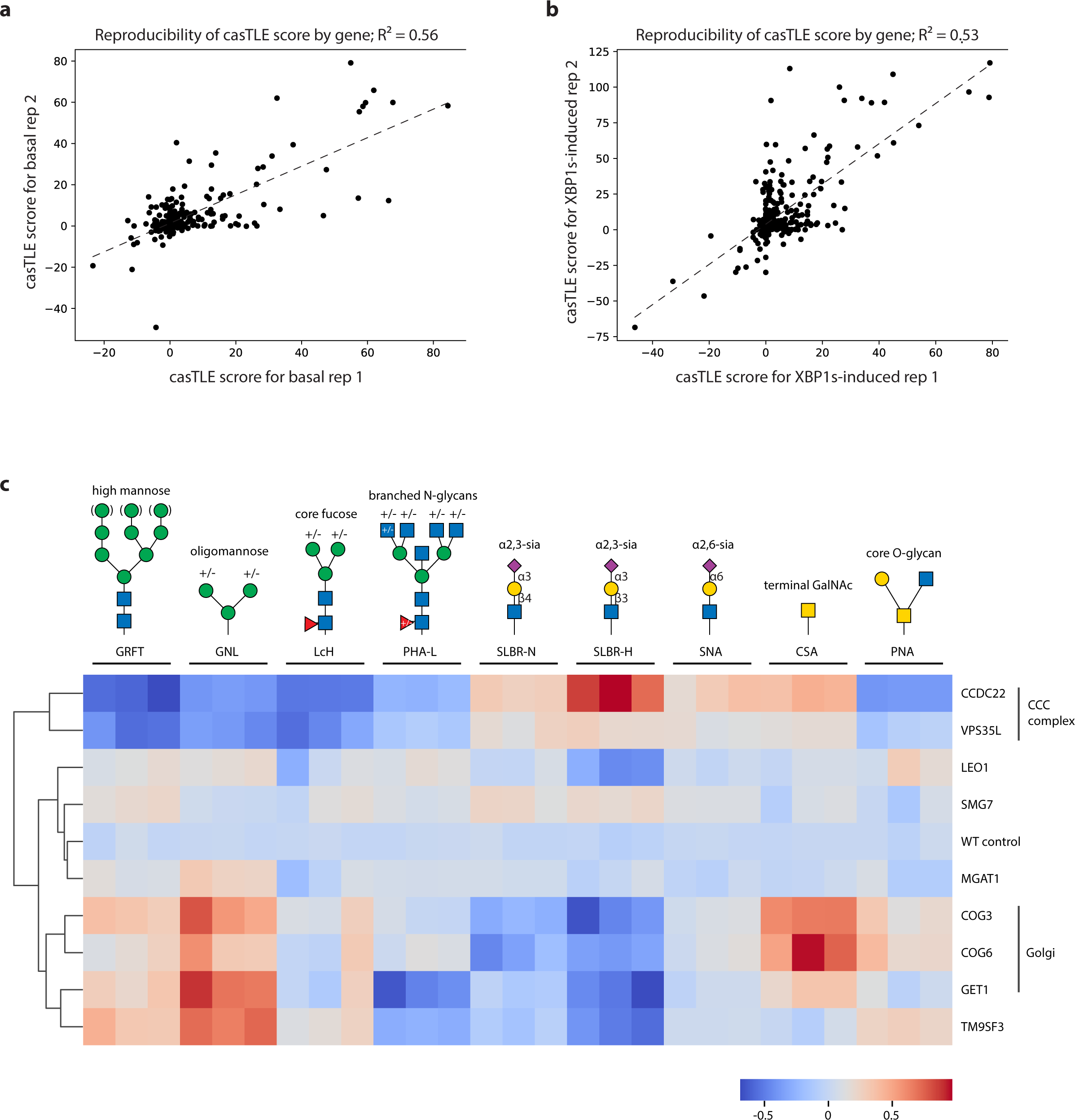
**a,** Reproducibility plot of casTLE scores for the CRISPRi targeted screen under basal condition. **b,** Reproducibility plot of casTLE scores for the CRISPRi targeted screen under XBP1s-indcued condition. **c,** Clustered heat map representing the results of competitive cell surface lectin binding assay for KD of top screen hits under basal conditions. Lectin binding specificities are indicated. Each column is a replicate and each row represents a different gene knock down.

**Extended Data Fig. 4.**
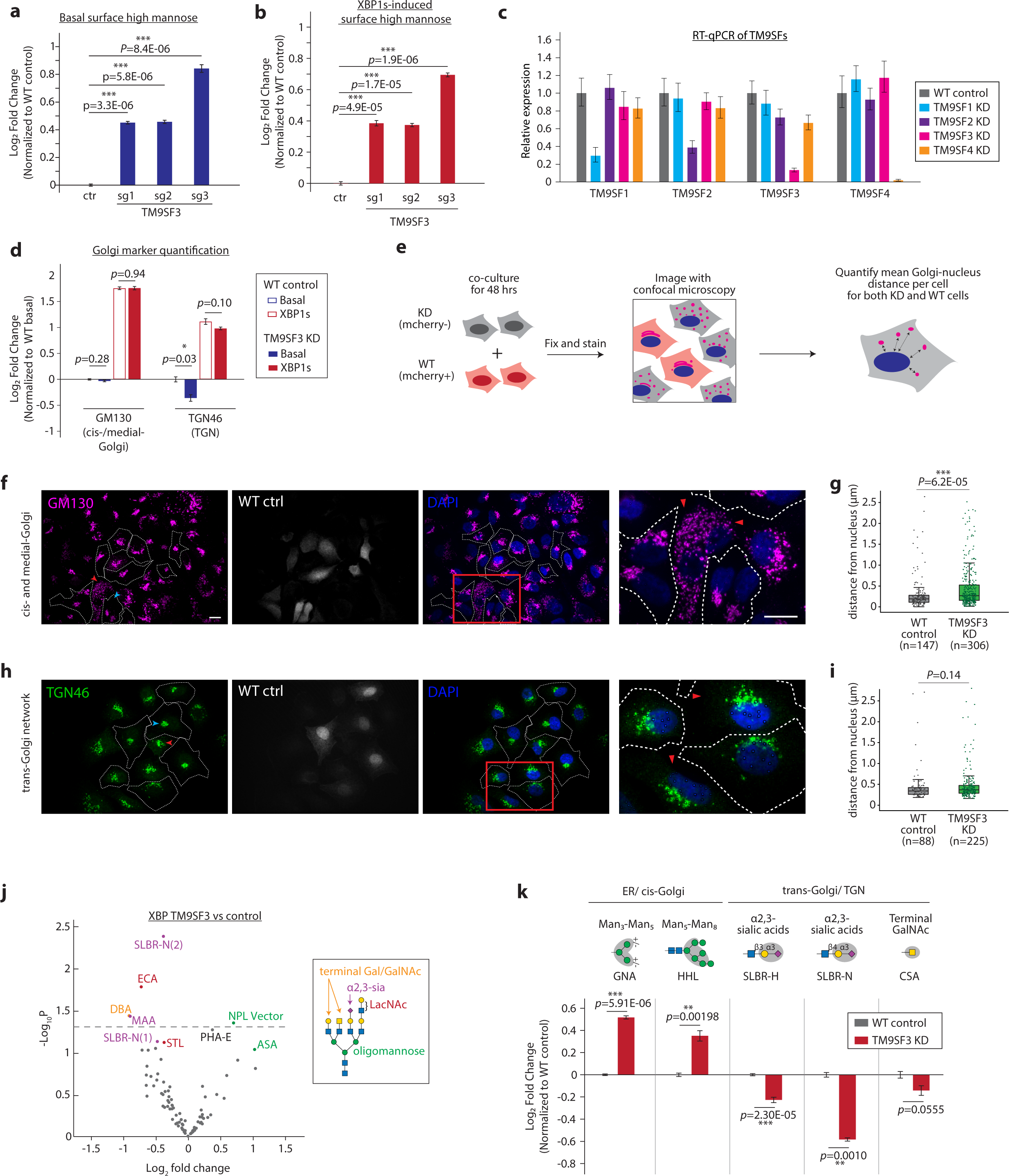
**a,** Competitive HHL binding assay in A549s with expressing three independent sgRNAs targeting TM9SF3. Data are presented as mean ± s.e.m. and are representative of two independent experiments performed in triplicate. **b,** Competitive HHL binding assay in A549s under basal conditions expressing three independent sgRNAs targeting TM9SF3. Data are presented as mean ± s.e.m. and are representative of two independent experiments performed in triplicate with consistent results. **c,** RT-qPCR for TM9SF family members. Gene expression is normalized to housekeeping genes GAPDH and HRPT1. **d,** Flow cytometry quantification of intracellular staining of GM130 and TGN46 in TM9SF3 knockdown cells under basal and XBP1s-induced conditions compared to wildtype control. Data are presented as mean ± s.e.m. and are representative of three independent experiments performed in triplicate with consistent results. **e,** Schematic for microscopy analysis. Wildtype control cells (mcherry-positive) were cocultured with TM9SF3 knockdown cells (mcherry-negtaive) as in-well internal controls. **f and h,** Representative confocal microscopy images of TM9SF3 knockdown and wildtype control cells, stained with cis-/medial-Golgi marker GM130 (f) or TGN marker TGN46 (h). Wildtype cells are outlined in dotted white lines. Magnified view of the red boxed areas are shown in the right-most column, with red arrows denoting KD cells. Scale bars, 10 µm. Images are representative of two independent experiments performed in triplicate. **g and i,** Quantification of average cis-/medial-Golgi or TGN distances from the nucleus in Extended Data Fig. 4f and 4h, respectively. Each data point represents the mean distance of a single cell. P-value is calculated by Mann-Whiteney U test. **j,** Volcano plot for lectin microarray results of XBP1s-induced A549 cells with TM9SF3 knocked down compared to wild type control. Lectins are color-coded by their glycan-binding specificities. **k,** Competitive cell surface lectin binding assay for TM9SF3 knocked down A549s compared to wildtype control under XBP1s-induced conditions. Lectin binding specificities and the location of where the modification predominately occurs are indicated. Data are presented as mean ± s.e.m. and are representative of two independent experiments performed in triplicate with consistent results.

**Extended Data Fig. 5.**
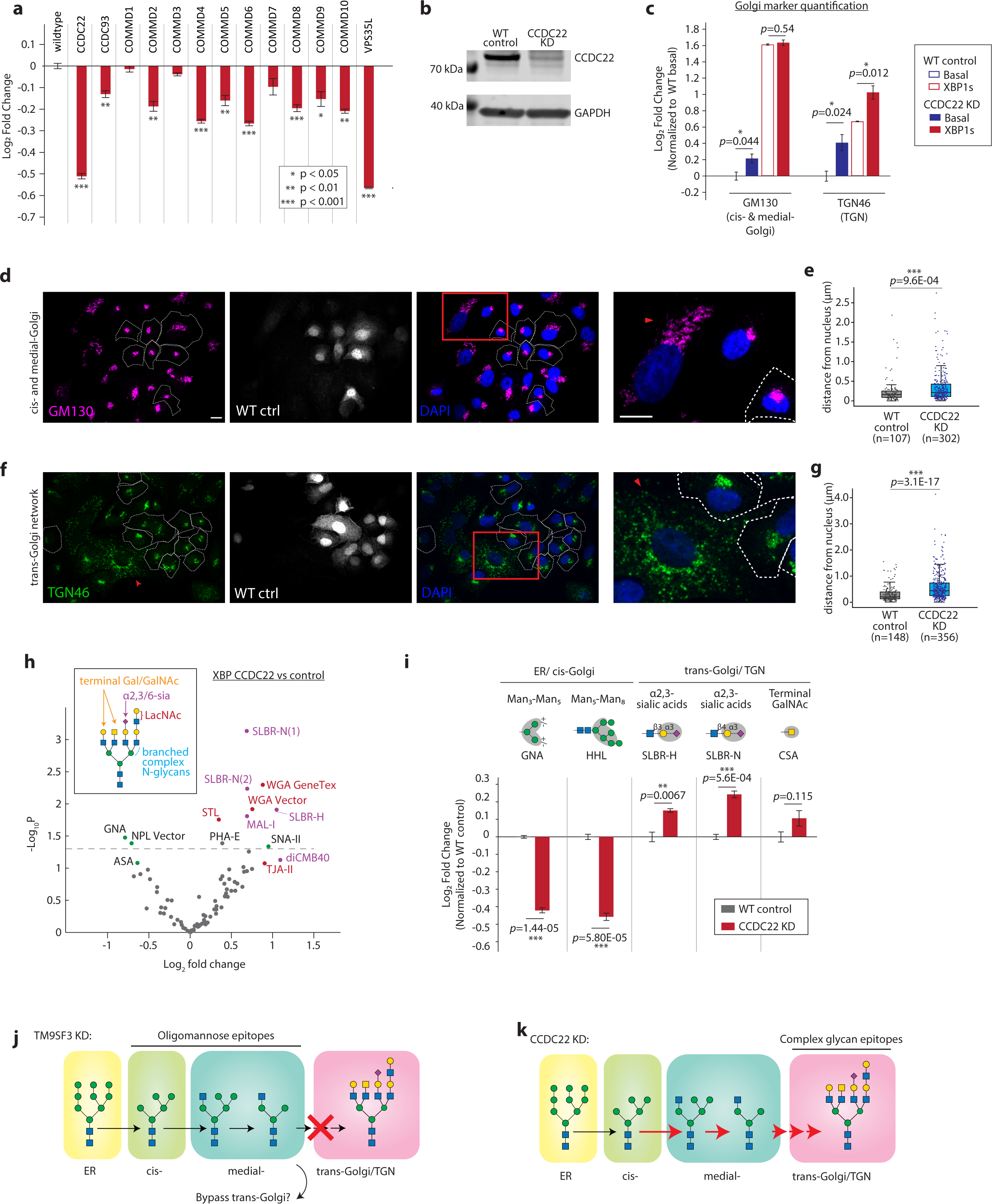
**a,** Competitive HHL binding assays on A549s under XBP1s-induced conditions for all knock down of all members of the CCC complex. Each gene is knocked down by co-expression of two independent sgRNAs. Data are presented as mean ± s.e.m. and are representative of three independent experiments performed in triplicate with consistent results. **b,** Western blot of CCDC22 knockdown cell line showing expected reduction in CCDC22 protein levels. **c,** Flow cytometry quantification of intracellular staining of GM130 and TGN46 in CCDC22 knockdown cells under basal and XBP1s-induced conditions compared to wildtype control. Data are presented as mean ± s.e.m. and are representative of three independent experiments performed in triplicate with consistent results. **d and f,** Representative confocal microscopy images of CCDC22 knockdown and wildtype control cells, stained with cis-/medial-Golgi marker GM130 (d) or TGN marker TGN46 (f). Wildtype cells are outlined in dotted white lines. Magnified view of the red boxed areas are shown in the right-most column, with red arrows denoting KD cells. Scale bars, 10 µm. Images are representative of two independent experiments performed in triplicate. **e and g,** Quantification of average cis-/medial-Golgi or TGN distances from the nucleus in Extended Data Fig. 5d and 5f, respectively. Each data point represents the mean distance of a single cell. P-value is calculated by Mann-Whiteney U test. **h,** Volcano plot for lectin microarray results of XBP1s-induced A549 cells with CCDC22 knocked down compared to wild type control. Lectins are color-coded by their glycan-binding specificities. **i,** Competitive cell surface lectin binding assay for CCDC22 knocked down A549s compared to wildtype control under XBP1s-induced conditions. Lectin binding specificities and the location of where the modification predominately occurs are indicated. Data are presented as mean ± s.e.m. and are representative of two independent experiments performed in triplicate with consistent results. **j,** Model for how glycosylation is altered in TM9SF3-KD cells: High mannose glycans are converted into oligomannose glycans in the cis- and medial-Golgi despite changes in morphology. However, the final stages of complex glycan synthesis, such as elongation by LacNAc motifs and capping with sialic acids, are inhibited. **k,** Model for how glycosylation is altered in CCDC22-KD cells: Despite the fragmented Golgi morphology, glycans are able to be remodeled from high mannose into complex glycans, perhaps at an enhanced efficiency. Resulting in a more complex N-glycan repertoire at the expense of high mannose glycans.

